# Temperate forest heterogeneity decreases local and landscape-scale spider diversity through habitat filtering despite species turnover

**DOI:** 10.1101/2025.08.29.673149

**Authors:** Jean-Léonard Stör, Julia Rothacher, Anne Chao, Maike Huszarik, Michael Junginger, Lisa Köstler-Albert, Oliver Mitesser, Akira S. Mori, Clara Wild, Jörg Müller

## Abstract

1. Spiders are key components of forest food webs, making use of the three-dimensional forest structure. Yet modern silviculture has homogenized temperate forest structure at local and landscape scales. The consequences of this homogenization for landscape-level spider diversity, however, remain largely unknown.
2. Therefore, we sampled spiders using pitfall traps across 234 patches in a large-scale, replicated field experiment at 11 paired European beech (*Fagus sylvatica)* forest sites in Germany. In one district per site, we experimentally diversified between-forest-patch complexity (ESBC) through canopy gap creation and deadwood enrichment and kept a second district untreated as a structurally homogeneous control.
3. We applied a novel meta-analytic framework to compare α-, β-, and γ-diversity of spiders between treatment and control districts, standardized for sample coverage, along Hill numbers giving increasing weight to abundance and for taxonomic, functional, and phylogenetic diversity. To gain a deeper insight into the effects of our intervention on the processes affecting the assembly of spider communities, we investigated the response of functional-phylogenetic diversity quantified by standardized effect sizes of mean pairwise distances (SES MFPD) to our treatments.
4. Based on 18,540 spider individuals from 206 species, treatment districts exhibited significantly lower γ- and α-diversity across all diversity facets and Hill numbers, particularly when focusing on rare species (q = 0). In contrast, β-diversity increased in treatment districts for phylogenetic and functional diversity across all Hill numbers (q = 0, 1, 2). Although spider abundances were higher in treatment patches, functional-phylogenetic diversity decreased in gaps and ESBC districts in general, indicating a shift from limiting similarity towards habitat filtering.
5. Our findings corroborate earlier results of high abundances but lower taxonomic and functional diversity in canopy gaps due to strong habitat filtering effects, resulting in overall lower α-diversity, which cannot be compensated by increasing β-diversity. Hence, the greater three-dimensional complexity of homogeneous forests supports more diverse spider metacommunities at the γ-scale, particularly when controlling for sample coverage, suggesting that canopy spider diversity in temperate forests may be underestimated. Nonetheless, higher abundances in treatment districts point to increased predator pressure and greater prey availability in structurally diversified forests.

## Introduction

A mosaic of connected, structurally diverse habitats is a key element for sustaining high biodiversity in landscapes (Müller et al., 2023; Schall et al., 2018; Tscharntke et al., 2012). Forest communities are shaped by natural disturbances, which diversify forests through accelerated tree senescence, but the effect range differs greatly for distinct organisms (Thorn et al., 2018; Viljur et al., 2022). Silvicultural practices in modern production forestry have led to a homogenization of the forest structure, reducing open woodland area throughout Europe and thereby limiting sun exposure and deadwood availability (Sing et al., 2018). The response of forest communities of early and late succession stages, which rely on a heterogenous landscape mosaic of frequent disturbances in the tree canopy, is strongly negative (Buckley, 2020). The reduction in structural complexity can be considered among the main causes of biotic homogenization on a local scale, but the effects of anthropogenic homogenization on diversity still remains unclear for numerous organismic groups (Keck et al., 2025).

Natural forests contain a wide array of different successional stages between early, mature and late ages with differing environmental conditions and species compositions (Hilmers et al., 2018). By conserving an array of complex woody elements, forests continue providing a structurally diverse habitat over changing forest generations (Uhl et al., 2022). Based on this, Keeton (2006) developed the concept of enhancing structural complexity (ESC) to restore forest heterogeneity. This approach increases variability in three-dimensional canopy density and retains different types and amounts of deadwood within forest stands. More recently, this concept has been further developed into measures for Enhancing Structural Beta-Complexity (ESBC) (Müller et al., 2023) which aim to create a diverse array of structurally complex habitats. Such structural enhancements promote the richness of numerous species groups, including plants, fungi, and animals (Heidrich et al., 2020) and deadwood diversity retention (Seibold et al., 2016).

A major group benefiting from deadwood and increased structural complexity are spiders (Graf et al., 2022; Müller et al., 2022). Spiders, a diverse group inhabiting ground and vegetation strata, are among the most abundant invertebrate predators and fulfill key ecological functions in virtually all terrestrial ecosystems (Michalko et al., 2019; Nyffeler & Birkhofer, 2017). Spiders also respond sensitively to changes in habitat structure, microclimatic conditions, and the availability of prey (Buchholz, 2016; Entling et al., 2007; Michalko & Birkhofer, 2021). In forests, ground-dwelling spiders are primarily influenced by changes in light availability and related factors, including canopy openness, understory cover and soil temperature (Hamřík et al., 2023; Samu et al., 2021). Accordingly, changes in environmental heterogeneity mediated by small-scale forestry treatments are reflected in the ground-dwelling spider β-diversity within forest stands (García-Tejero et al., 2018; Plath et al., 2024). Spider abundance and diversity on α-level has been shown to be mitigated mainly by structural heterogeneity (Heidrich et al., 2020; Plath et al., 2025), canopy cover (Müller et al., 2022; Vierling et al., 2011) and anthropogenic disturbances (Plath et al., 2021). Both spider abundance and species numbers are often higher in early successional habitats than in old-growth stands. However, these studies have primarily focused on local (α) diversity, while the effects of forest structural change on β- and γ-diversity remain poorly understood. β-diversity, defined as the variability in assemblage composition among forest patches, may increase with environmental heterogeneity and, in turn, elevate γ-diversity. This highlights the potential role of ESBC measures in shaping spider diversity at the landscape scale.

Yet, some obstacles exist when assessing diversity. When sampling highly diverse taxa, such as arthropods, it is inevitable that some species remain undetected. Consequently, spider samples are always an incomplete subset of the local community (Chao, Gotelli, et al., 2014). Sample coverage quantifies this completeness, defined as the proportion of individuals in the community belonging to detected species (Chao & Jost, 2012). Coverage has been shown to vary across categories in spider pitfall-trap samples from forest reserves (Huber et al., 2025) and to be consistently lower in open-canopy patches for other arthropods than spiders (Rothacher et al., 2025). Without correcting for such biases, diversity can be under- or overestimated, and in the worst case, apparent treatment effects may simply reflect differences in sample coverage. Since readily available tools enable accounting for these sampling effects (Chao et al., 2021; Hsieh et al., 2016), coverage-based standardization using extrapolation and rarefaction is increasingly used for unbiased biodiversity assessments (Kortmann, Chao, Chiu, et al., 2025; Rothacher et al., 2025).

In this study, we made use of one of the largest temperate forest manipulation experiments in Europe (Müller et al., 2023) to assess how forest heterogeneity impacts spider diversity. In this paired experimental design, silvicultural manipulations were conducted in formerly homogeneous broadleaved production forests in Germany. These manipulations created gradients in canopy openness and increased the structural variety of deadwood, both important features of natural old-growth forests (Bauhus et al., 2009; Donato et al., 2012). This allowed us to test the effects of heterogeneous vs. homogenous forests on spider diversity standardized for sample coverage across multiple spatial scales. Specifically, we hypothesized that: (i) structural heterogeneity enhances spider diversity at both the patch scale (α-diversity) and the landscape scale (γ-diversity) compared to structurally homogeneous control forests; (ii) increased structural heterogeneity promotes the assembly of distinct spider assemblages across patches, resulting in higher between-patch β-diversity in heterogeneous forests.

## Material and Methods

### Study area and experimental design

This study is part of the collaborative research unit ΒETA-FOR (Müller et al., 2023) and was conducted in 11 experimental forest sites across Germany, all dominated by European beech (*Fagus sylvatica*) with smaller, site-specific proportions (1–30% of total basal area) of other broad-leaved and coniferous species. The forest sites are located in the management zone of the Bavarian Forest National Park (4 sites: B04, B05, B06, B07), the University Forest of the Julius-Maximilians-Universität Würzburg (3 sites: U01, U02, U03), the Hunsrück-Hochwald National Park (1 site: H09) and the administrative districts of Passau (1 site: P08), Saarland (1 site: S10) and Lübeck (1 site: L11) spanning a broad elevational gradient from 38 m (Lübeck) to 1143 m a.s.l. (Bavarian Forest) (Fig. 1). Consequently, the experimental design captures a wide range of macroclimatic conditions, soil types, and tree species compositions representative of Central European beech forest types, enabling broadly applicable insights into how forest structure influences biodiversity in these ecosystems.

**Figure 1.**
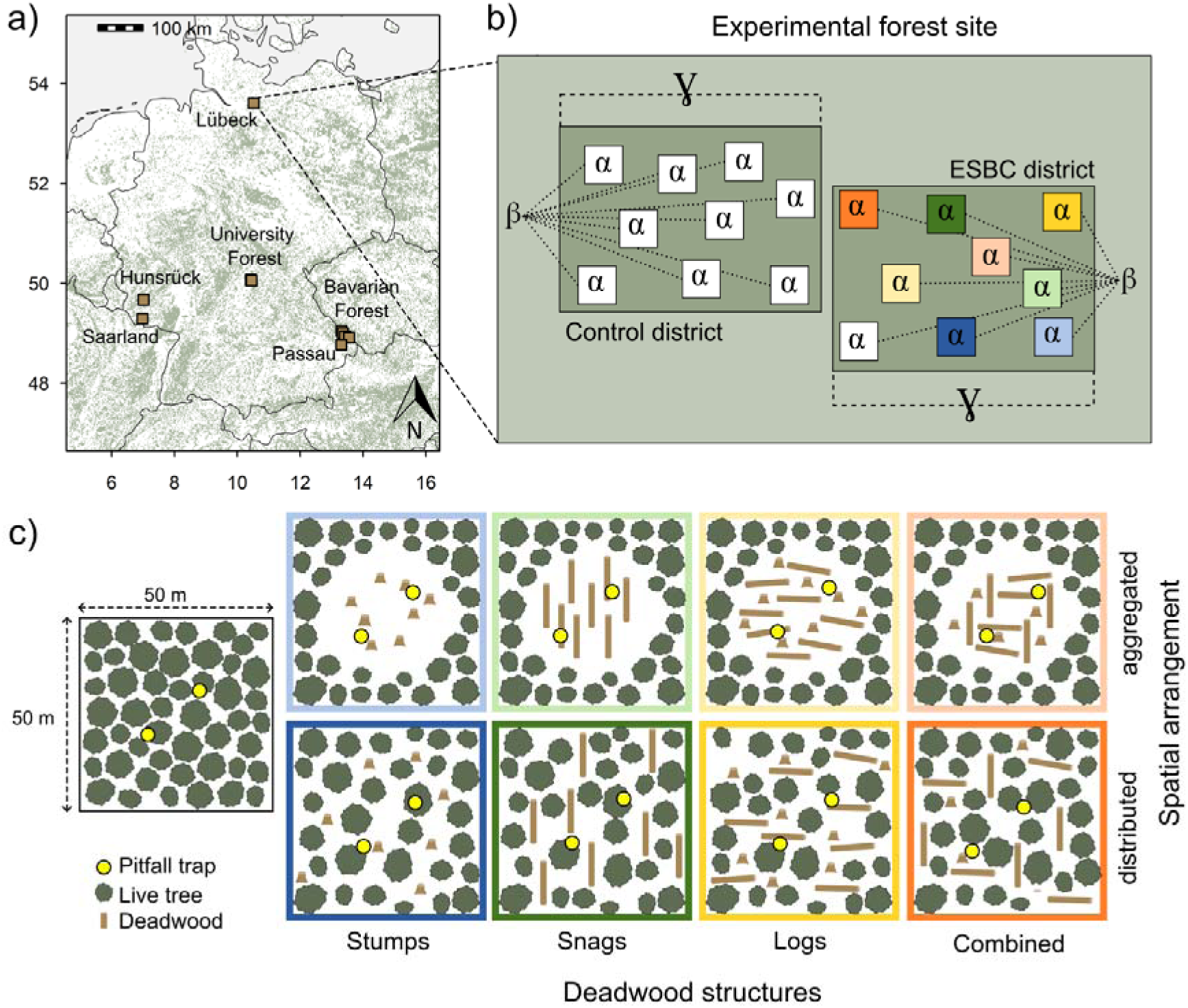
Experimental design of the ΒETA-FOR research unit. (a) Location of the eleven experimental forest sites across six regions in Germany indicated by brown rectangles; forest cover is shown in green. Note that three sites are located in the University Forest and four in the Bavarian Forest. (b) Schematic representation of an experimental forest site, consisting of a structurally heterogeneous treatment (ESBC) district and a structurally homogeneous control district. (c) Overview of the nine treatment types implemented in each ESBC district. The position of pitfall traps in each patch are indicated by yellow dots. Note that in the University Forest, an additional number of six distinct treatment types per district and landscape were applied. For an illustration of the full set of treatment types, see Figure S1.

In each site, two experimental districts were established, each consisting of a fixed set of 50 m × 50 m patches (Fig. 1). Sites in the Bavarian Forest, Passau, Saarland, Lübeck, and Hunsrück each comprised 18 patches (9 patches per district), while sites in the University Forest contained 30 patches (15 patches per district). In total, this design resulted in 234 experimental patches across Germany.

In one district per site (treatment district), we experimentally enhanced the structural beta complexity (ESBC) between patches through targeted silvicultural interventions. These were implemented in the winters of 2015/16 (Bavarian Forest, Passau), 2016/17 (Saarland, Lübeck, Hunsrück), and 2018/19 (University Forest). Interventions were applied either spatially aggregated by creating canopy gaps (∼30 m in diameter) in the patch center or distributed throughout the patch and maintaining a closed canopy. In both spatial settings, interventions created different deadwood structures (i.e., stumps, snags, logs, and a combination thereof) resulting in each deadwood type occurring in both sunny (canopy gap) and shady (no gap) conditions. These manipulations resulted in nine distinct treatment types in each landscape (see Fig. 1). In the University Forest, an additional six treatment types were applied (i.e., stump extraction, habitat tree creation, and retention of tree crowns on site in both sunny and shady conditions; see Fig. S1 for details).

The control district was established by selecting the same number of patches as in the corresponding ESBC district, located in close proximity to it. To ensure comparability, each control district shared similar elevation and tree species composition with its associated ESBC district. Since no interventions were applied, the control patches closely resemble structurally homogeneous production forests, as commonly found across large areas of Central Europe. They are characterized by mature trees of similar age in the early to mid-optimum stage of stand development, a closed canopy, and a limited amount of deadwood. This landscape scale experimental approach enables assessing within-patch α-diversity, between-patch β-diversity, and overall γ-diversity within each district of a landscape (Fig. 1).

## Spider Sampling

Spiders were sampled using two pitfall traps per patch. The pitfall traps were installed in the central area of each patch (25 × 25 m) to represent the patch conditions well (Fig. 1). Although primarily designed for ground-dwelling arthropods, pitfall traps are also able to sample vegetation dwelling spiders, as some descend from the canopy during the mating season (Müller et al., 2022). Each trap consisted of a 400 ml plastic cup buried at ground level. We covered each trap with a transparent plastic roof (15 × 15 cm, approximately 5 cm above the ground), to protect the trap from rain or leaves. We used a solution of sodium chloride (NaCl) as the trapping fluid and added a drop of odorless detergent to break the surface tension, which ensured rapid killing of the sampled arthropods. The traps were active for three consecutive months each year (i.e. May, June, July) and emptied once per month. Patches in the Bavarian Forest, University Forest and Passau were sampled in 2022, patches in Saarland, Lübeck, Hunsrück were sampled in 2023 with permits obtained from the respective regional authorities (Regierung von Niederbayern; Regierung von Unterfranken; Ministerium für Umwelt, Klima, Mobilität, Agrar und Verbraucherschutz Saarland; Landesamt für Umwelt Schleswig-Holstein). The collected samples were sorted and spiders from each sample were identified to species level following the taxonomy of Roberts (1985).

## Data preparation

Data preparation and statistical analyses were performed using R 4.2.2 (R Core Team, 2025). In a first step, we restricted our dataset to adult spiders and aggregated the data on patch level, by summing up the abundances from all spider individuals per patch. To assess phylogenetic diversity, we created a phylogenetic tree encompassing all spider species of our dataset (Fig. 2). We used the spider phylogeny by Müller et al. (2022) as a backbone and integrated 13 spider species additionally present in our data based on their sister taxon or close relatives using the *bind.tip* function of the *phytools* package (Revell, 2024).

**Figure 2.**
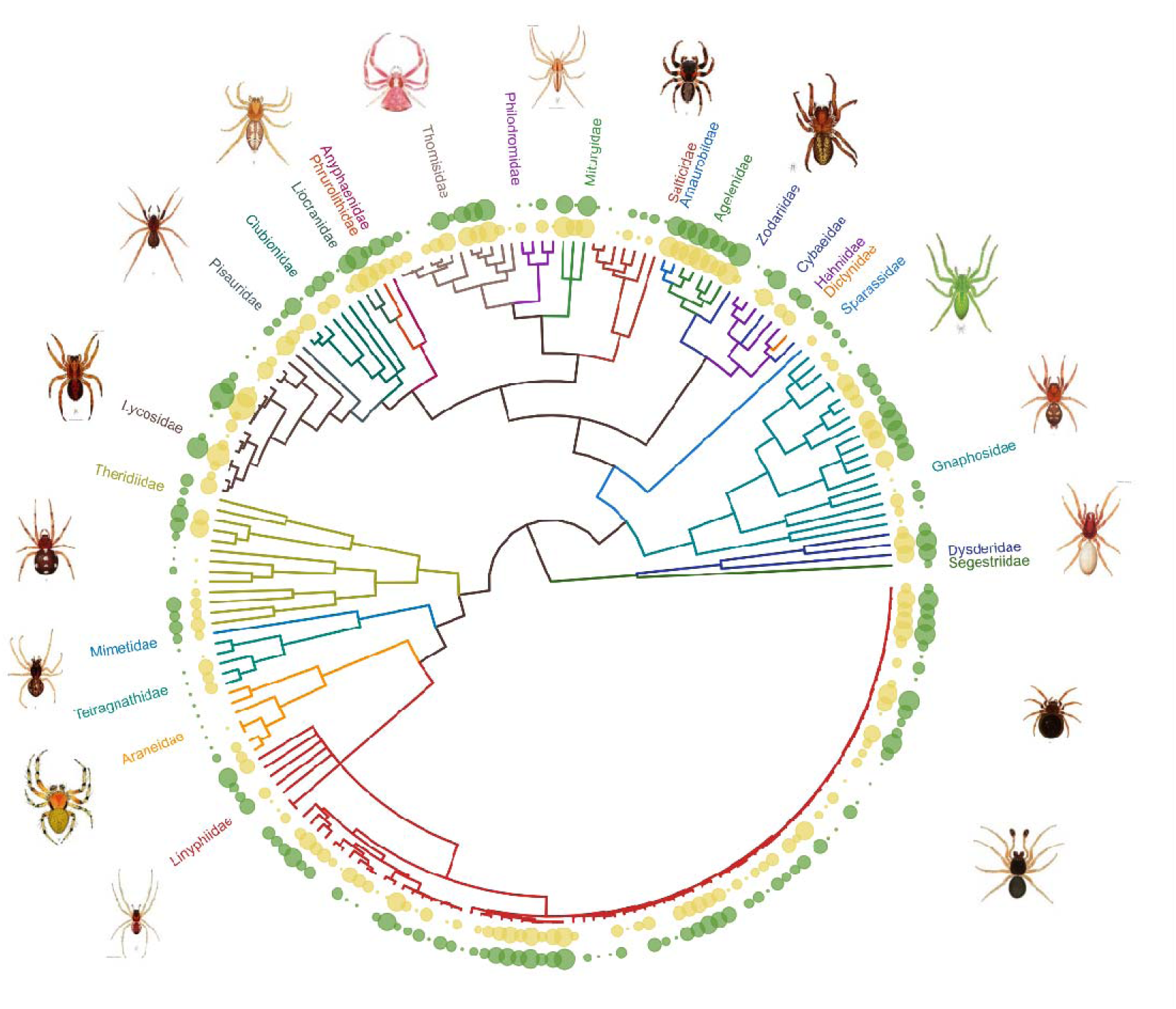
Phylogenetic tree of 206 spider species sampled within the forest manipulation experiment demonstrates the dominance of sheet weavers (Linyphiidae) in temperate forests. Each tip of the phylogeny represents one species, family association indicated by colors. Dot size at the tree tips refers to the abundance of each species across treatments: inside (yellow) ESBC, outside (green) Control. Illustrations of spider families adapted from Roberts, 1985.

We further gathered information on spider functional traits, by integrating 63 additional spider species to the trait dataset of Müller et al. (2022). This dataset comprises ecological traits, including hunting strategy, preferred vertical stratum, ballooning ability, and mean body size, together with a set of morphological trait measurements (Table 1). For spider species absent from the original dataset, ecological traits were added from the literature, and morphological traits were measured directly. All morphological traits were standardized to body length by extracting residuals from a linear regression of log_e_(trait) ∼ log_e_(body length). Functional distance between species was calculated using Gower distance (see R code; Bässler et al 2014, Cadotte et al 2013).

**Table 1.** Traits characterizing the behavior of spider species in response to local environments.

| Trait | Ecology | Measurement | Source |
| --- | --- | --- | --- |
| Stratum | Spiders use different strata in a forest from ground to canopy layer | 1/0 coding of 5 forest strata: epigeic, herb layer, shrub layer, tree trunk, canopy layer; multiple strata per species possible | Müller et al. (2023) and additional information from literature |
| Web-type | Spiders use different types of web for foraging | Five categories: no web, web with tangles of stopping threads, horizontal web bottom, horizontal web top, vertical web | Müller et al. (2023) and additional information from literature |
| Balloonig | Some spiders use passive ballooning for long distance dispersal | 1/0 coding of the ballooning high and ballooning low probability | Birkhofer, Meub, et al. (2015) and additional information from literature |
| Body size | Body size decrease from warm and dry to cool and moist conditions, body size increase with habitat complexity | Continuous value from literature | Entling et al (2010) and additional information from literature |
| Opisthosoma and prosoma breadth | Arthropods are narrower in structurally more complex habitats | Residuals of a linear model $\log_e(y) \sim \log_e(x)$: Opisthosoma and prosoma breadth (mm) ~ body length (mm) measured | Gibb et al. (2015) and additional information from literature |
| Leg length | Might decrease in structurally simple habitats | Residuals of tibia length of first leg as before | Gibb et al. (2015) and additional information from literature |
| Fang length | Fast prey in less complex and warmer environment evokes long fang length | Residuals of chelicera length as before | Gibb et al. (2015) and additional information from literature |

## Statistical analysis

To evaluate taxonomic (TD), phylogenetic (PD) and functional diversity (FD) differences between ESBC and Control districts, we employed a comparative meta-analysis approach that integrates the two key concepts sample coverage and Hill numbers within the iNEXT.3D framework (Chao et al., 2021), thus providing robust and comparable biodiversity estimates across sites. To evaluate the landscape-scale effects of our experimental manipulation on spider γ-diversity, we conducted the meta-analysis across the 11 paired experimental districts using the *iNEXTmeta* function (Chao, 2025) for abundance data with 99 bootstrapping replications. Because diversity data are typically incomplete, it is inevitable for meaningful comparisons to standardize to a common level of sample coverage, which can vary substantially in open- and closed-canopy forests. The *iNEXTmeta* function incorporates sample coverage standardization and the calculation of diversity metrics along Hill numbers (q = 0, 1, 2) to consider species frequency distributions (Hill, 1973). This allows a focus on rare (q = 0), common (q = 1), and dominant (q = 2) species and lineages in the dataset. In this analysis TD quantifies the effective number of equally abundant species. PD quantifies the effective total branch length and FD quantifies the effective number of functional species. Since TD, PD and FD are in the same units (species/lineage equivalents) they can be meaningfully compared (Chao, Chiu, et al., 2014). All samples were standardized to sample coverage of 0.76, which represents the default coverage provided by the *iNEXTmeta* function. Ultimately, *iNEXTmeta* provides average effect sizes and bootstrap-derived confidence intervals aggregated across district pairs.

To identify the relevant spatial scale of treatment effects we decomposed γ-diversity (district level) into two hierarchical components: (1) within-patch diversity (α component), and (2) among-patch diversity (β component) using the *iNEXTmeta_beta* function (Chao, 2025) within the meta-analysis framework. The overall effect was considered significant when the 95% confidence interval did not include zero. The functions are currently available at: https://github.com/AnneChao/iNEXT.meta. For TD, γ-diversity quantifies the effective number of species in the pooled community across all patches, while α-diversity represents the average effective number of species per patch. The interpretation for PD and FD follows analogously by replacing “species” with “lineages” or “functional units”. β-diversity is defined as the ratio of γ-diversity to α-diversity. In a district with nine patches, β-diversity equals 1 when all patches share identical species composition and abundances and reaches a maximum of 9 when no species are shared among patches, indicating complete turnover. Thus, β-diversity quantifies the degree of compositional differentiation among patches and is expressed as the effective number of distinct patches. To enable meaningful comparisons of β-diversity while accounting for variation in patch numbers across districts (i.e., 9 or 15 patches per district), we used the Jaccard-type turnover (1-S) transformation (Chao et al., 2019), which quantifies dissimilarity between assemblages relative to γ-diversity and is standardized by patch number, with zero indicating identical assemblages in the patches and one indicating no shared species.

To gain deeper insights into the effects of our treatment on local, patch- and district-wise assembly processes we calculated standardized effects sizes of mean pairwise distances. As in many trait datasets important information is missing, we first followed the approach by Cadotte et al. (2013). In this approach ecological dissimilarity is calculated by dissimilarity in traits (functional dissimilarity) with stepwise adding information from dissimilarity in phylogenies as surrogate for unmeasured traits. Here we used the same phylogenetic and functional distances (FD and PD) distances between species as in the meta-analyses above. Based on these distance matrices, we calculated the standardized effect size of the mean pairwise distance between co-occurring species within assemblages of a patch abundance weighted. The effect size was calculated based on null models with 999 randomizations by reshuffling the tip labels to achieve independence from species numbers (Webb et al., 2002). Standardized effect value above 1.96 indicate limitations in niche overlap, while values below -1.96 indicate environmental filtering (Cadotte et al., 2013; Müller et al., 2022). Adding stepwise phylogenetic distance to the functional distance revealed the highest R² for a-value of 12% phylogenetic information in the between species distance (see Fig. S11), which was used as the final value for extracting the standardized effect size of mean pairwise functional-phylogenetic diversity (SES MFPD) for all patches (for detailed description see Supplementary Text). This SES MFPD as well as the abundance and the number of observed species per patch was finally modelled with generalized additive models using the *gam* function from the *mgcv* package (Wood, 2011). We used spider abundance, observed number of species, and SES MFPD values as response variables, and included treatment (ESBC vs. control), canopy (open vs. closed), and deadwood (enriched, i.e. treatments with snags, logs, habitat trees, or crown deadwood, vs. non-enriched) as explanatory variables. Forest site was included as a smoothed non-linear spline to account for repeated measurements and the spatial dependence of patches within the same site.

## Results

We analyzed a total of 18,540 spiders belonging to 206 species which represents approximately 20% of all spider species occurring in Germany (Blick et al., 2016). Across Hill numbers, taxonomic, functional, and phylogenetic γ- and α-diversity were significantly lower in ESBC districts compared to control districts (Fig. 3a, b; Fig. S2-4, Fig. S8-10). This effect was strongest for taxonomic diversity, suggesting that many species are functionally or phylogenetically redundant. The reduction was most pronounced for rare (q = 0) compared to common (q = 1) and dominant (q = 2) species, lineages, and functional units. Taxonomic β-diversity was higher for rare species (q = 0) but decreased with diversity order (Fig. 3c; Fig. S5-7). Phylogenetic and functional β-diversity in contrast showed an increase with diversity order and highest values for dominant functional species or phylogenetic lineages (q = 2) (Fig. 3c, Fig. S5-7).

**Figure 3.**
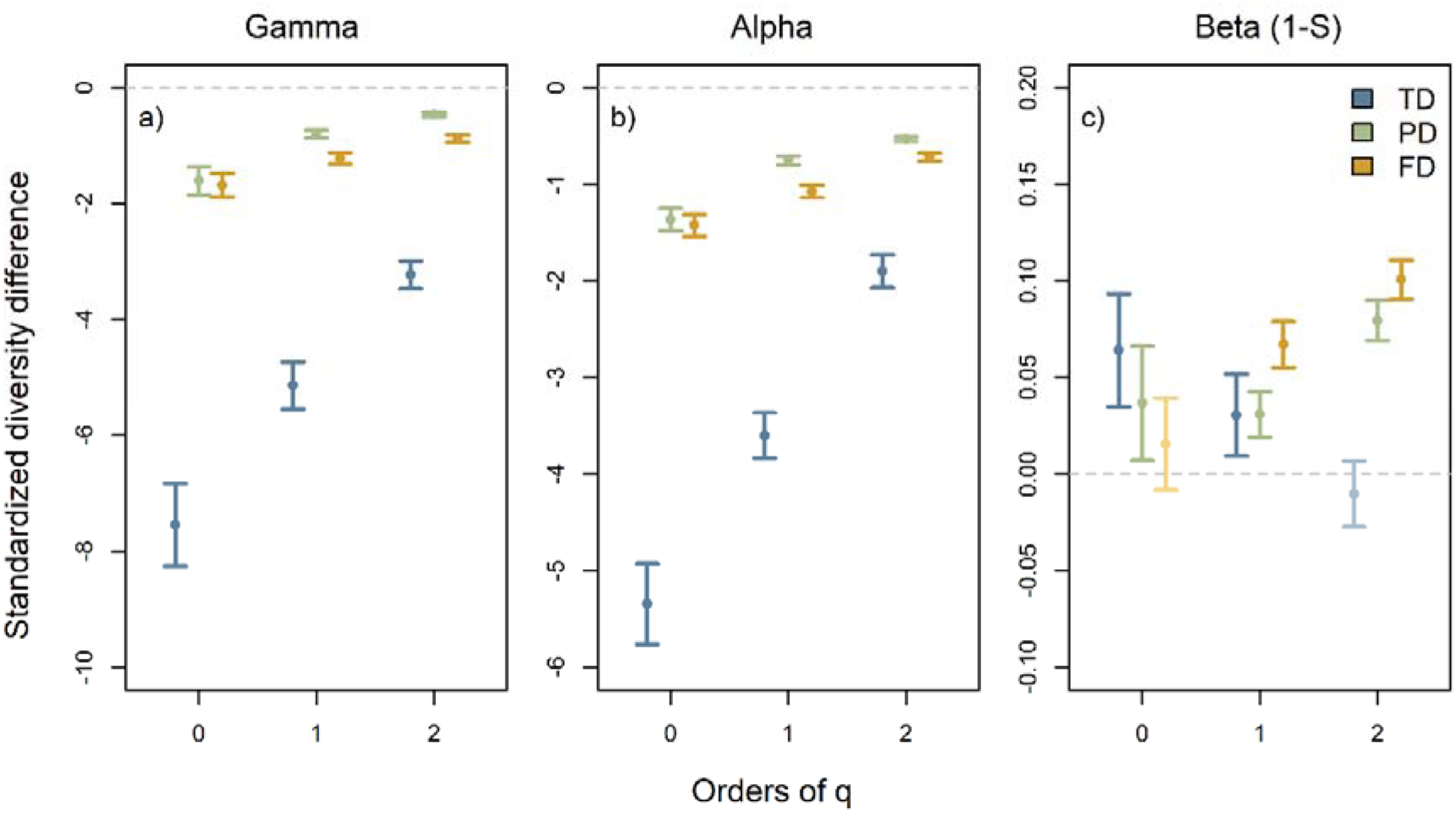
Meta-analysis of the spider diversity response across 11 experimental forest sites. Results are shown for landscape-scale γ-diversity (a), patch-scale α-diversity (b) and multiplicative (1 − S) β-diversity (c). Points represent predicted means; error bars denote 95% confidence intervals at a fixed coverage level of 0.76. Standardized diversity differences are calculated between each pair of control and ESBC districts in directly comparable species-equivalent units for γ- and α-diversity, and for taxonomic (TD, blue) and phylogenetic (PD, green) and functional diversity (FD, yellow) across three diversity orders (q = 0, 1, 2). Positive differences indicate an increase in diversity from control to structurally enhanced districts, while negative differences indicate a diversity decrease. Significant effects are shown in dark shades, while non-significant results (confidence intervals include zero) are shown in light shades of blue, green and yellow.

A detailed analysis of the patch-level assembly processes revealed that canopy openness and ESBC treatments increased spider abundance, whilst deadwood enrichment reduced abundance and observed species number (Tab. 2). Functional diversity declined with canopy openness and ESBC but remained unaffected by deadwood (Tab. 2). Standardized effect sizes of mean pairwise functional distances were strongly negative under canopy opening and ESBC, indicating environmental clustering of spider communities.

**Table 2.**
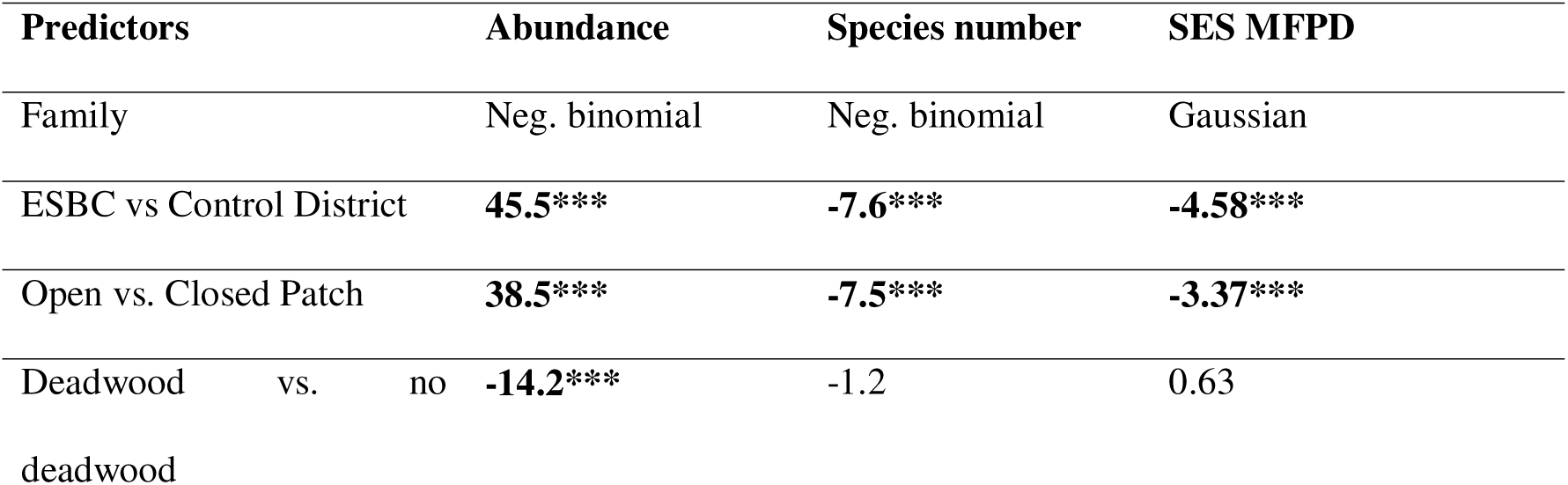
Z-scores from generalized additive models testing the effects of ESBC establishment, canopy (open vs. closed), and deadwood enrichment on spider abundance, observed species number, and standardized effect sized of the mean pairwise functional-phylogenetic distance (SES MFPD, a-value 12%) . Significant results are shown in bold. Significance levels: *p < 0.05; **p < 0.01; ***p < 0.001.

## Discussion

Contrary to our first hypothesis, structural heterogenization reduced spider α- and γ-diversity compared to control forests, although β-diversity partly increased in line with our second hypothesis. Spider abundance was higher in heterogeneous stands and open canopies, but these assemblages were composed of functionally more similar species.

### Habitat filtering and standardized sampling explain spider diversity loss in heterogeneous landscapes

Our large-scale forest manipulation experiment demonstrated that experimental heterogenization of beech forests can reduce spider diversity by up to eight species at the landscape scale, alongside their associated functions. Similar reductions in spider diversity were observed at the local patch-scale. Open forest structures provide an increased amount of understory vegetation (Bradler et al., 2025), whilst the complex three-dimensional structure of tree canopies is reduced. Consequently, our small-scale open-canopy patches offered microhabitats typical of early- and mid-successional stages, resulting in distinct species assemblages, as also reported after forest management alterations (Hamřík et al., 2023, 2024; Plath et al., 2024; Viljur et al., 2022). Thus, ESBC patches were colonized by a variety of species, which ecologically complemented and, in some cases, even replaced closed-forest species (see Fig. 2). Similar results were observed in Müller et al. (2022), where small-scale canopy gaps also hosted a functionally different set of spiders.

In line with (García et al., 2014; Tsang et al., 2025), the partly-higher spider β-diversity in our study was unable to outweigh the lower local α-diversity in the ESBC districts through species turnover. This was ultimately reflected in lower patch-scale α- and landscape scale γ-diversity. The increase in β-diversity that fails to offset a decline in α-diversity has recently been demonstrated for fragmented habitats (Gonçalves-Souza et al., 2025). Even total γ-diversity was lower in fragmented compared to homogeneous forests, largely driven by stronger negative α-effects. This highlights the necessity of examining biodiversity at multiple scales rather than focusing solely on α-diversity, as is most commonly done. In our experiment, spiders proved to be rather unique in their responses: while beetles showed higher α- and β-diversity under ESBC treatments and consequently higher γ-diversity (Mitesser et al., 2025), plants primarily exhibited pronounced α-increases without corresponding β-effects, a pattern also observed in hoverflies (Bradler et al., 2025; Massó Estaje et al., 2025). Together with recent meta-analyses (Gonçalves-Souza et al., 2025; Keck et al., 2025; Viljur et al., 2022) our systematic experiment further highlights the importance of assessing biodiversity responses to human pressures across spatial scales and taxonomic groups. At first glance, previous studies appear to contrast with our results by reporting higher species numbers in open habitat patches (Košulič et al., 2016; Plath et al., 2025). However, these comparisons were often based on raw species counts (Chao & Jost, 2012; Gotelli & Colwell, 2001), often misinterpreted as richness, without accounting for sample coverage. When standardized to a common level of sample coverage, in contrast, post-disturbance management scenarios did not affect spider species richness in spruce forests (Plath et al., 2024). In our study, open-canopy patches were dominated by cursorial species, which are more likely to be captured in pitfall traps, leading to higher local abundance and more complete sampling. Observed sample coverage (Fig. S11) was higher in ESBC patches with gaps. Thus, observed richness in the ESBC closely approximates true richness, whereas failure to standardize for this bias underestimates spider diversity, particularly in closed-canopy forests. Whenever systematic biases in sample coverage arise along ecological gradients or experimental treatments, comparisons of observed diversity become inappropriate (Kortmann, Chao, Chiu, et al., 2025; Kortmann, Chao, Schaefer, et al., 2025; Lettenmaier et al., 2022; Rothacher et al., 2025).

Our data show that canopy spiders form highly diverse communities that use the full three-dimensional forest space, but this diversity is strongly reduced in canopy gaps. An earlier fogging study at one of our sites already revealed high canopy diversity and stochastic recolonization dynamics after winter losses (Hsieh & Linsenmair, 2011). These processes result in less environmentally filtered communities in closed canopies (Hsieh, 2011), a pattern confirmed in our large-scale replicated comparison of gaps and intact stands (Müller et al., 2022). Given the high dispersal capacity and reproductive ability of spiders (Müller et al., 2022; Schmidt & Tscharntke, 2005), incomplete colonization of ESBC patches can be ruled out. Meta-community analyses indicate that ballooning enables broad dispersal (Entling et al., 2007; Samu et al., 2018), allowing species to respond to local disturbances such as our ESBC patches. Species replacement was therefore driven primarily by dispersal and habitat filtering, consistent with findings from other studies reshaping local communities studies (Carvalho et al., 2020; Plath et al., 2024; Samu et al., 2021). In fine-grained patchiness, such as single ESBCs within intact forest stands, edge effects and spillover did not override species responses to altered environmental conditions (Frost et al., 2015). Consequently, spider diversity on ESBC patches was lower because remnant forest species inhabiting canopies, tree trunks, or the cool, moist forest floor were reduced and replaced by new settlers benefiting from increased light, structural diversity, and variable humidity (García-Tejero et al., 2018; Hamřík et al., 2023).

### Functional and phylogenetic diversity responses indicate ecological similarity

We consistently observed the strongest effects of ESBC on taxonomic spider diversity, followed by phylogenetic and functional responses. This pattern suggests that many taxonomic species are subsumed within phylogenetic and functional units, indicating a high degree of ecological similarity in spiders, where multiple species perform similar ecological functions. Functional similarity, or niche overlap, can act as an insurance mechanism against environmental fluctuations, buffering communities against species loss so that ecosystem process rates remain stable (Eisenhauer et al., 2023). To date, little is known about the rates of species versus trait loss and turnover in invertebrate communities facing land-use change (Rigal et al., 2018). However, in spiders, functional loss and decreased niche overlap have been linked to species loss and anthropogenic disturbance in tropical forests (Potapov et al., 2020). Similarly, silvicultural interventions in the form of ESBC treatments led to a reduction in functional diversity of spiders of the whole district. Our results on the assembly patterns with reduced SES MFPD further underlines that gaps act as strong habitat filter for species with similar traits, reducing the overall functional diversity in ESBC. Similar filtering effects have been shown for spiders in forests gaps of different size (Müller et al., 2022).

### Sampling spider assemblages with pitfall traps comes with limitations

Spiders occupy the three-dimensional space of a forest, including tree canopies. Although we detected some arboreal spiders in our pitfall trap samples, it is important to consider the limitations of this method when interpreting our results. Pitfall traps are biased towards large-sized and high-mobility taxa (e.g., ground-hunting, running, stalking spiders) on the ground level, where inhabitants of the canopy and shrub layer rarely venture (Turnbull, 1973). Thus, canopy-dwelling species are generally underrepresented in our passive ground-level trapping system. Yet, in line with Müller et al. (2022) we were able to capture a number of tree-dwelling spider species like *Tetragnatha pinicola* (Tetragnathidae)*, Cyclosa conica* (Araneidae) or *Diaea dorsata* (Thomisidae) which thrive on leaves and branches, as well as camouflaged trunk-hunters like *Clubiona caerulescens* (Clubionidae), *Philodromus aureolus* (Philodromidae) and *Segestria senoculata* (Segestriidae) with pitfall traps in stands with a closed canopy. This is because temperate forest spiders regularly descend from the canopy during the mating season, which coincides with our sampling period in spring and early summer. Thus, the responses of rare (q = 0) spiders are partly driven by these species and their functional and phylogenetic traits, highlighting the advantage of using Hill numbers in the analysis. On open-canopy patches, also in line with Müller et al. (2022), we were not able to capture these canopy-dwelling species. This suggests that they either move shorter distances when on the ground or simply occur at lower densities in areas with fewer trees, reducing the likelihood of being incidentally captured. For more comprehensive sampling and inclusion of the diverse canopy fauna, future studies should employ targeted combinations of methods across forest strata. This could include canopy fogging (Hsieh, 2011; Müller et al., 2018), vegetation sampling using beating trays (Klimm et al., 2025), and pitfall trapping for epigeic species (Müller et al., 2022), providing more complete insights into spider diversity responses in temperate forests.

## Conclusion

Our replicated forest heterogenization experiment provided novel insights into how homogeneous versus heterogeneous forest structures shape spider diversity across scales. At the local patch scale, our results corroborate previous findings that forest openings harbor high spider abundances but reduced taxonomic and functional diversity due to strong habitat filtering. At the landscape scale, however, β-diversity increased in line with the habitat heterogeneity hypothesis. Yet, this turnover did not offset the loss of α-diversity in canopy gaps, ultimately reducing γ-diversity. This scale-dependent reduction in γ-diversity appears unusual compared to other taxa, highlighting that biodiversity responses to structural heterogenization can vary both across spatial scales and taxonomic groups.

## Acknowledgement

We thank all local managers and technical staff for their support in our research. We are particularly grateful to the interns of the Bavarian Forest National Park for field work and technical support. This research was supported by funding from the German Research Foundation (DFG) within the research unit ‘ΒETA-FOR’ (project no. 459717468), with additional support from the Julius-Maximilians-Universität Würzburg (JMU). JR acknowledges additional funding from the Bavarian State Ministry for Food, Agriculture, Forestry and Tourism (StMELF) (grant no. L062).

## Conflict of Interest

The authors declare that they have no known competing financial interests or personal relationships that could have appeared to influence the work reported in this paper.

## Author Contributions

Jörg Müller conceived the ideas and designed the methodology; Julia Rothacher, Clara Wild, Jean-Léonard Stör, Michael Junginger and Lisa Köstler-Albert collected the data; Julia Rothacher, Jörg Müller and Jean-Léonard Stör analyzed the data; Anne Chao and Oliver Mitesser provided resources for the analysis; Julia Rothacher visualized the results; Jean-Léonard Stör and Julia Rothacher led the writing of the manuscript; Akira S. Mori and Maike Huszarik supported the writing of the manuscript with their comments. All authors gave final approval for publication.

## Notes

https://zenodo.org/records/14766448

### Competing Interest Statement

The authors have declared no competing interest.

## References

Bauhus, J., Puettmann, K., & Messier, C. (2009). Silviculture for old-growth attributes. Forest Ecology and Management, 258, 525–537. 10.1016/j.foreco.2009.01.053

Blick, T., Finch, O., Harms, K., Kiechle, J., Kielhorn, K., Kreuels, M., Malten, A., Martin, D., Muster, C., Nährig, D., Platen, R., Rödel, I., Scheidler, M., Staudt, A., Stumpf, H., & Tolke, D. (2016). Rote Liste und Gesamtartenliste der Spinnen (Arachnida: Araneae) Deutschlands. Naturschutz Und Biologische Vielfalt, 70(4), 383. – 510.

Bradler, P. M., Delory, B. M., Dittrich, S., Cadotte, M. W., Chao, A., Härdtle, W., Mitesser, O., Mori, A., Müller, J., Nishizawa, K., Plas, F. van der, Oheimb, G. von, & Fichtner., A. (2025). Structurally heterogeneous forest landscapes promote gamma diversity, but not beta diversity, in forest understorey plant communities. Www.Biorxiv.Org, submitted.

Buchholz, S. (2016). Natural peat bog remnants promote distinct spider assemblages and habitat specific traits. Ecological Indicators, 60, 774–780. 10.1016/j.ecolind.2015.08.025

Buckley, P. (2020). Coppice restoration and conservation: a European perspective. Journal of Forest Research, 25(3), 125–133. 10.1080/13416979.2020.1763554

Cadotte, M., Albert, C. H., & Walker, S. C. (2013). The ecology of differences: assessing community assembly with trait and evolutionary distances. Ecology Letters, 16(10), 1234–1244. 10.1111/ele.12161

Cadotte, M., & Davies, J. (2016). Phylogenies in Ecology: A Guide to Concepts and Methods. 10.1515/9781400881192

Carvalho, J. C., Malumbres-Olarte, J., Arnedo, M. A., Crespo, L. C., Domenech, M., & Cardoso, P. (2020). Taxonomic divergence and functional convergence in Iberian spider forest communities: Insights from beta diversity partitioning. Journal of Biogeography, 47(1), 288–300. 10.1111/jbi.13722

Chao, A. (2025). CRAN repository iNEXT functions. https://github.com/AnneChao/iNEXT.meta

Chao, A., Chiu, C.-H., & Jost, L. (2014). Unifying Species Diversity, Phylogenetic Diversity, Functional Diversity, and Related Similarity and Differentiation Measures Through Hill Numbers. Annual Review of Ecology, Evolution, and Systematics, 45, 297–324. 10.1146/annurev-ecolsys-120213-091540

Chao, A., Chiu, C., Wu, S.-H., Huang, C.-L., & Lin, Y. (2019). Comparing two classes of alpha diversities and their corresponding beta and (dis)similarity measures, with an application to the Formosan sika deer (Cervus nippon taiouanus) reintroduction program. Methods in Ecology and Evolution, 10. 10.1111/2041-210x.13233

Chao, A., Gotelli, N. J., Hsieh, T. C., Sander, E. L., Ma, K. H., Colwell, R. K., & Ellison, A. M. (2014). Rarefaction and extrapolation with Hill numbers: a framework for sampling and estimation in species diversity studies. Ecological Monographs, 84(1), 45–67. 10.1890/13-0133.1

Chao, A., Henderson, P. A., Chiu, C.-H., Moyes, F., Hu, K.-H., Dornelas, M., & Magurran, A. E. (2021). Measuring temporal change in alpha diversity: A framework integrating taxonomic, phylogenetic and functional diversity and the iNEXT.3D standardization. Methods in Ecology and Evolution, 12(10), 1926–1940. 10.1111/2041-210X.13682

Chao, A., & Jost, L. (2012). Coverage-based rarefaction and extrapolation: standardizing samples by completeness rather than size. Ecology, 93(12), 2533–2547. 10.1890/11-1952.1

Donato, D., Campbell, J., & Franklin, J. (2012). Multiple successional pathways and precocity in forest development: can some forests be born complex? Journal of Vegetation Science, 23, 576–584. 10.2307/23251088

Eisenhauer, N., Hines, J., Maestre, F. T., & Rillig, M. C. (2023). Reconsidering functional redundancy in biodiversity research. Npj Biodiversity, 2(1), 9. 10.1038/s44185-023-00015-5

Entling, W., Entling, M., Bacher, S., Brandl, R., & Nentwig, W. (2007). Niche properties of Central European spiders: Shading, moisture and the evolution of the habitat niche. Global Ecology and Biogeography, 16, 440–448. 10.1111/j.1466-8238.2006.00305.x

Frost, C. M., Didham, R. K., Rand, T. A., Peralta, G., & Tylianakis, J. M. (2015). Community-level net spillover of natural enemies from managed to natural forest. Ecology, 96(1), 193–202. 10.1890/14-0696.1

García-Tejero, S., Spence, J. R., O’Halloran, J., Bourassa, S., & Oxbrough, A. (2018). Natural succession and clearcutting as drivers of environmental heterogeneity and beta diversity in North American boreal forests. PLoS ONE, 13(11), 1–16. 10.1371/journal.pone.0206931

García, D., Martínez, D., Stouffer, D. B., & Tylianakis, J. M. (2014). Exotic birds increase generalization and compensate for native bird decline in plant–frugivore assemblages. Journal of Animal Ecology, 83(6), 1441–1450. 10.1111/1365-2656.12237

Gonçalves-Souza, T., Chase, J. M., Haddad, N. M., Vancine, M. H., Didham, R. K., Melo, F. L. P., Aizen, M. A., Bernard, E., Chiarello, A. G., Faria, D., Gibb, H., de Lima, M. G., Magnago, L. F. S., Mariano-Neto, E., Nogueira, A. A., Nemésio, A., Passamani, M., Pinho, B. X., Rocha-Santos, L., … Sanders, N. J. (2025). Species turnover does not rescue biodiversity in fragmented landscapes. Nature, 640(8059), 702–706. 10.1038/s41586-025-08688-7

Gotelli, N. J., & Colwell, R. K. (2001). Quantifying biodiversity: procedures and pitfalls in the measurement and comparison of species richness. Ecology Letters, 4(4), 379–391. 10.1046/j.1461-0248.2001.00230.x

Graf, M., Seibold, S., Gossner, M. M., Hagge, J., Weiß, I., Bässler, C., & Müller, J. (2022). Coverage based diversity estimates of facultative saproxylic species highlight the importance of deadwood for biodiversity. Forest Ecology and Management, 517, 120275. 10.1016/j.foreco.2022.120275

Hamřík, T., Gallé, R., & Košulič, O. (2024). Ecologically sustainable retention forestry supports spider biodiversity in the Lower Morava UNESCO Biosphere Reserve. Insect Conservation and Diversity. 10.1111/icad.12765

Hamřík, T., Košulič, O., Gallé, R., Gallé-Szpisjak, N., & Hédl, R. (2023). Opening the canopy to restore spider biodiversity in protected oakwoods. Forest Ecology and Management, 541, 121064. 10.1016/j.foreco.2023.121064

Heidrich, L., Bae, S., Levick, S., Seibold, S., Weisser, W., Krzystek, P., Magdon, P., Nauss, T., Schall, P., Serebryanyk, A., Wöllauer, S., Ammer, C., Bässler, C., Doerfler, I., Fischer, M., Gossner, M. M., Heurich, M., Hothorn, T., Jung, K., … Müller, J. (2020). Heterogeneity–diversity relationships differ between and within trophic levels in temperate forests. Nature Ecology and Evolution, 4(9), 1204–1212. 10.1038/s41559-020-1245-z

Hill, M. O. (1973). Diversity and Evenness: A Unifying Notation and Its Consequences. Ecology, 54(2), 427–432. 10.2307/1934352

Hilmers, T., Friess, N., Bässler, C., Heurich, M., Brandl, R., Pretzsch, H., Seidl, R., & Müller, J. (2018). Biodiversity along temperate forest succession. Journal of Applied Ecology, 55(6), 2756–2766. 10.1111/1365-2664.13238

Hsieh, T. C., Ma, K. H., & Chao, A. (2016). iNEXT: an R package for rarefaction and extrapolation of species diversity (Hill numbers). Methods in Ecology and Evolution, 7(12), 1451–1456. 10.1111/2041-210X.12613

Hsieh, Y.-L., & Linsenmair, K. E. (2011). Underestimated spider diversity in a temperate beech forest. Biodiversity and Conservation, 20, 2953–2965. 10.1007/s10531-011-0158-1

Hsieh, Y. L. (2011). The diversity and ecology of the spider communities of European beech canopy.

Huber, A., Fahrig, L., Seidl, R., Erhardt, A. T., Müller, J., & Seibold, S. (2025). Inferring the role of habitat heterogeneity in SLOSS (single large or several small) for beetles, spiders, and birds in forest reserves. Biological Conservation, 311, 111403. 10.1016/j.biocon.2025.111403

Keck, F., Peller, T., Alther, R., Barouillet, C., Blackman, R., Capo, E., Chonova, T., Couton, M., Fehlinger, L., Kirschner, D., Knüsel, M., Muneret, L., Oester, R., Tapolczai, K., Zhang, H., & Altermatt, F. (2025). The global human impact on biodiversity. Nature, 641(8062), 395–400. 10.1038/s41586-025-08752-2

Keeton, W. S. (2006). Managing for late-successional/old-growth characteristics in northern hardwood-conifer forests. Forest Ecology and Management, 235(1), 129–142. 10.1016/j.foreco.2006.08.005

Kembel, S. W., Cowan, P. D., Helmus, M. R., Cornwell, W. K., Morlon, H., Ackerly, D. D., Blomberg, S. P., & Webb, C. O. (2010). Picante: R tools for integrating phylogenies and ecology. Bioinformatics, 26(11), 1463–1464. 10.1093/bioinformatics/btq166

Klimm, F. S., Boetzl, F. A., König, S., Bräu, M., Burtchen, L., Mandery, K., Stör, J.-L., & Krauss, J. (2025). Life at the (h)edge—Multidiversity in shrub ecotones is driven by habitat quality and shrub foliage cover. Journal of Applied Ecology, 62(6), 1520–1530. 10.1111/1365-2664.70061

Kortmann, M., Chao, A., Chiu, C.-H., Heibl, C., Mitesser, O., Moriniere, J., Bozicevic, V., Hothorn, T., Rothacher, J., Englmeier, J., Ewald, J., Fricke, U., Ganuza, C., Haensel, M., Moning, C., Redlich, S., Rojas Botero, S., Tobisch, C., Uhler, J., & Müller, J. (2025). A shortcut to sample coverage standardization in metabarcoding data provides new insights into land-use effects on insect diversity. Proceedings B, 292. 10.1098/rspb.2024.2927

Kortmann, M., Chao, A., Schaefer, H. M., Blüthgen, N., Gelis, R., Tremlett, C. J., Busse, A., Püls, M., Seibold, S., Kriegel, P., Rabl, D., de la Hoz, M., Şekercioğlu, Ç. H., Schleuning, M., Feldhaar, H., Newell, F. L., Kümmet, S., Mitesser, O., Peters, M. K., & Müller, J. (2025). Sample coverage affects diversity measures of bird communities along a natural recovery gradient of abandoned agriculture in tropical lowland forests. Journal of Applied Ecology, 62(3), 480–491. 10.1111/1365-2664.14879

Košulič, O., Michalko, R., & Hula, V. (2016). Impact of canopy openness on spider communities: Implications for conservation management of formerly coppiced oak forests. PLoS One, 11(2), e0148585. 10.1371/journal.pone.0148585

Lettenmaier, L., Seibold, S., Bässler, C., Brandl, R., Gruppe, A., Müller, J., & Hagge, J. (2022). Beetle diversity is higher in sunny forests due to higher microclimatic heterogeneity in deadwood. Oecologia, 198. 10.1007/s00442-022-05141-8

Massó Estaje, C., Rothacher, J., Vujić, A., Miličić, M., Chao, A., Mitesser, O., Müller, J., Claßen, A., & Steffan-Dewenter, I. (2025). Experimental enhancement of structural heterogeneity in forest landscapes promotes multidimensional hoverfly diversity. BioRxiv, 2025.08.22.670311. 10.1101/2025.08.22.670311

Michalko, R., & Birkhofer, K. (2021). Habitat niches suggest that non-crop habitat types differ in quality as source habitats for Central European agrobiont spiders. Agriculture, Ecosystems & Environment, 308, 107248. 10.1016/j.agee.2020.107248

Michalko, R., Pekár, S., Dul’a, M., & Entling, M. H. (2019). Global patterns in the biocontrol efficacy of spiders: A meta-analysis. Global Ecology and Biogeography, 28(9), 1366– 1378. 10.1111/geb.12927

Mitesser, O., Cadotte, M., Akira Mori, Plas, F. van der, Chao, A., Rothacher, J., Bässler, C., Bevanda, M., Biedermann, P., Bradler, P., Castaneda-Gomez, A., Decker, O., Delory, B. M., Dittrich, S., Fichtner, A., Kreis, A., Köstler-Albert, L., von Oheimb, G., Pflumm, L., … Müller, J. (2025). Old growth attributes by chain saw: how between-patch heterogeneity changes the metacommunities of beetles in temperate forests. Ecology Letters, submitted.

Müller, J., Brandl, R., Brändle, M., Förster, B., de Araujo, B. C., Gossner, M. M., Ladas, A., Wagner, M., Maraun, M., Schall, P., Schmidt, S., Heurich, M., Thorn, S., & Seibold, S. (2018). LiDAR-derived canopy structure supports the more-individuals hypothesis for arthropod diversity in temperate forests. Oikos, 127(6), 814–824. 10.1111/oik.04972

Müller, J., Brandl, R., Cadotte, M. W., Heibl, C., Bässler, C., Weiß, I., Birkhofer, K., Thorn, S., & Seibold, S. (2022). A replicated study on the response of spider assemblages to regional and local processes. Ecological Monographs, 92(3), 1–19. 10.1002/ecm.1511

Müller, J., Mitesser, O., Cadotte, M. W., van der Plas, F., Mori, A. S., Ammer, C., Chao, A., Scherer-Lorenzen, M., Baldrian, P., Bässler, C., Biedermann, P., Cesarz, S., Claßen, A., Delory, B. M., Feldhaar, H., Fichtner, A., Hothorn, T., Kuenzer, C., Peters, M. K., … Eisenhauer, N. (2023). Enhancing the structural diversity between forest patches—A concept and real-world experiment to study biodiversity, multifunctionality and forest resilience across spatial scales. Global Change Biology, 29(6), 1437–1450. 10.1111/gcb.16564

Nyffeler, M., & Birkhofer, K. (2017). An estimated 400-800 million tons of prey are annually killed by the global spider community. Science of Nature, 104(3–4). 10.1007/s00114-017-1440-1

Pearse, W. D., Cadotte, M. W., Cavender-Bares, J., Ives, A. R., Tucker, C. M., Walker, S. C., & Helmus, M. R. (2015). pez: phylogenetics for the environmental sciences. Bioinformatics, 31(17), 2888–2890. 10.1093/bioinformatics/btv277

Plath, E., Böhme, W., Fischer, D., Griebel, L., Jochims, K., Schreek, K., Thiem, C., & Fischer, K. (2025). Spider diversity in a disturbed forest landscape highlights the importance of management heterogeneity. Insect Conservation and Diversity, 18(3), 400–416. 10.1111/icad.12815

Plath, E., Fischer, D., Glebsattel, S., Naeckel, L., & Fischer, K. (2024). Salvage Logging and Secondary Succession Promote Spider Diversity in Post-disturbance Stands in Western Germany. Forest Science, 70. 10.1093/forsci/fxae014

Plath, E., Rischen, T., Mohr, T., & Fischer, K. (2021). Biodiversity in agricultural landscapes: Grassy field margins and semi-natural fragments both foster spider diversity and body size. Agriculture, Ecosystems & Environment, 316, 107457. 10.1016/j.agee.2021.107457

Potapov, A. M., Dupérré, N., Jochum, M., Dreczko, K., Klarner, B., Barnes, A. D., Krashevska, V., Rembold, K., Kreft, H., Brose, U., Widyastuti, R., Harms, D., & Scheu, S. (2020). Functional losses in ground spider communities due to habitat structure degradation under tropical land-use change. Ecology, 101(3), e02957. 10.1002/ecy.2957

Revell, L. J. (2024). phytools 2.0: an updated R ecosystem for phylogenetic comparative methods (and other things). PeerJ, 12, e16505. 10.7717/peerj.16505

Rigal, F., Cardoso, P., Lobo, J. M., Triantis, K. A., Whittaker, R. J., Amorim, I. R., & Borges, P. A. V. (2018). Functional traits of indigenous and exotic ground-dwelling arthropods show contrasting responses to land-use change in an oceanic island, Terceira, Azores. Diversity and Distributions, 24(1), 36–47. 10.1111/ddi.12655

Roberts, M. J. (1985). Die Spinnen von Grossbritannien und Irland (3rd ed.).

Rothacher, J., Seidl, R., Thom, D., Kortmann, M., Chao, A., Chiu, C.-H., Heibl, C., Hothorn, T., Mitesser, O., Mori, A. S., Morinière, J., Pierick, K., Wild, C., Wild, N., & Müller, J. (2025). The impact of tree mortality and post-disturbance management on insect diversity in temperate forests: Insights from a replicated experiment. Journal of Applied Ecology, 62(8), 1878–1888. 10.1111/1365-2664.70086

Samu, F., Elek, Z., Kovács, B., Fülöp, D., Botos, E., Schmera, D., Aszalós, R., Bidló, A., Németh, C., Sass, V., Tinya, F., & Ódor, P. (2021). Resilience of spider communities affected by a range of silvicultural treatments in a temperate deciduous forest stand. Scientific Reports, 11(1), 1–13. 10.1038/s41598-021-99884-8

Samu, F., Horváth, A., Neidert, D., Botos, E., & Szita, É. (2018). Metacommunities of spiders in grassland habitat fragments of an agricultural landscape. Basic and Applied Ecology, 31. 10.1016/j.baae.2018.07.009

Schall, P., Gossner, M. M., Heinrichs, S., Fischer, M., Boch, S., Prati, D., Jung, K., Baumgartner, V., Blaser, S., Böhm, S., Buscot, F., Daniel, R., Goldmann, K., Kaiser, K., Kahl, T., Lange, M., Müller, J., Overmann, J., Renner, S. C., … Ammer, C. (2018). The impact of even-aged and uneven-aged forest management on regional biodiversity of multiple taxa in European beech forests. Journal of Applied Ecology, 55(1), 267–278. 10.1111/1365-2664.12950

Schmidt, M. H., & Tscharntke, T. (2005). Landscape context of sheetweb spider (Araneae: Linyphiidae) abundance in cereal fields. Journal of Biogeography, 32(3), 467–473. 10.1111/j.1365-2699.2004.01244.x

Seibold, S., Bässler, C., Brandl, R., Büche, B., Szallies, A., Thorn, S., Ulyshen, M. D., & Müller, J. (2016). Microclimate and habitat heterogeneity as the major drivers of beetle diversity in dead wood. Journal of Applied Ecology, 53(3), 934–943. 10.1111/1365-2664.12607

Sing, L., Metzger, M. J., Paterson, J. S., & Ray, D. (2018). A review of the effects of forest management intensity on ecosystem services for northern European temperate forests with a focus on the UK. Forestry, 91(2), 151–164. 10.1093/forestry/cpx042

Team, R.-C. (2022). R: A Language and Environment for Statistical Computing. https://www.r-project.org/

Thorn, S., Bässler, C., Brandl, R., Burton, P. J., Cahall, R., Campbell, J. L., Castro, J., Choi, C.-Y., Cobb, T., Donato, D. C., Durska, E., Fontaine, J. B., Gauthier, S., Hebert, C., Hothorn, T., Hutto, R. L., Lee, E.-J., Leverkus, A. B., Lindenmayer, D. B., … Müller, J. (2018). Impacts of salvage logging on biodiversity: A meta-analysis. Journal of Applied Ecology, 55(1), 279–289. 10.1111/1365-2664.12945

Tsang, T. P. N., De Santis, A. A. A., Armas-Quiñonez, G., Ascher, J. S., Ávila-Gómez, E. S., Báldi, A., Ballare, K. M., Balzan, M. V, Banaszak-Cibicka, W., Bänsch, S., Basset, Y., Bates, A. J., Baumann, J. M., Beal-Neves, M., Bennett, A., Bezerra, A. D. M., Blochtein, B., Bommarco, R., Brosi, B., … Bonebrake, T. C. (2025). Land Use Change Consistently Reduces α-But Not β- and γ-Diversity of Bees. Global Change Biology, 31(1), e70006. 10.1111/gcb.70006

Tscharntke, T., Tylianakis, J., Rand, T., Didham, R., Fahrig, L., Batary, P., Bengtsson, J., Clough, Y., Crist, T., Dormann, C., Ewers, R., Fründ, J., Holt, R., Holzschuh, A., Klein, A., Kleijn, D., Kremen, C., Landis, D., Laurance, W., & Westphal, C. (2012). Tscharntke et al 2012 Landscape moderation of biodiversity patterns and processes - eight hypotheses.

Turnbull, A. L. (1973). Ecology of the true spiders (Araneomorphae). Annual Review of Entomology, 18, 305–348.

Uhl, B., Krah, F.-S., Baldrian, P., Brandl, R., Hagge, J., Müller, J., Thorn, S., Vojtech, T., & Bässler, C. (2022). Snags, logs, stumps, and microclimate as tools optimizing deadwood enrichment for forest biodiversity. Biological Conservation, 270, 109569. 10.1016/j.biocon.2022.109569

Vierling, K. T., Bässler, C., Brandl, R., Vierling, L. A., Weiß, I., & Müller, J. (2011). Spinning a laser web: predicting spider distributions using LiDAR. Ecological Applications, 21(2), 577–588. 10.1890/09-2155.1

Viljur, M.-L., Abella, S. R., Adámek, M., Alencar, J. B. R., Barber, N. A., Beudert, B., Burkle, L. A., Cagnolo, L., Campos, B. R., Chao, A., Chergui, B., Choi, C.-Y., Cleary, D. F. R., Davis, T. S., Dechnik-Vázquez, Y. A., Downing, W. M., Fuentes-Ramirez, A., Gandhi, K. J. K., Gehring, C., … Thorn, S. (2022). The effect of natural disturbances on forest biodiversity: an ecological synthesis. Biological Reviews, 97(5), 1930–1947. 10.1111/brv.12876

Webb, C., Ackerly, D., Mcpeek, M., & Donoghue, M. (2002). Phylogenies and Community Ecology. Annu. Rev. Ecol. Syst, 8, 475–505. 10.1146/annurev.ecolsys.33.010802.150448

Wood, S. N. (2011). Fast Stable Restricted Maximum Likelihood and Marginal Likelihood Estimation of Semiparametric Generalized Linear Models. Journal of the Royal Statistical Society Series B: Statistical Methodology, 73(1), 3–36. 10.1111/j.1467-9868.2010.00749.x

