## Supplementary material for "Temperate forest heterogeneity decreases local and landscape-scale spider diversity through habitat filtering despite species turnover": Fig. S1

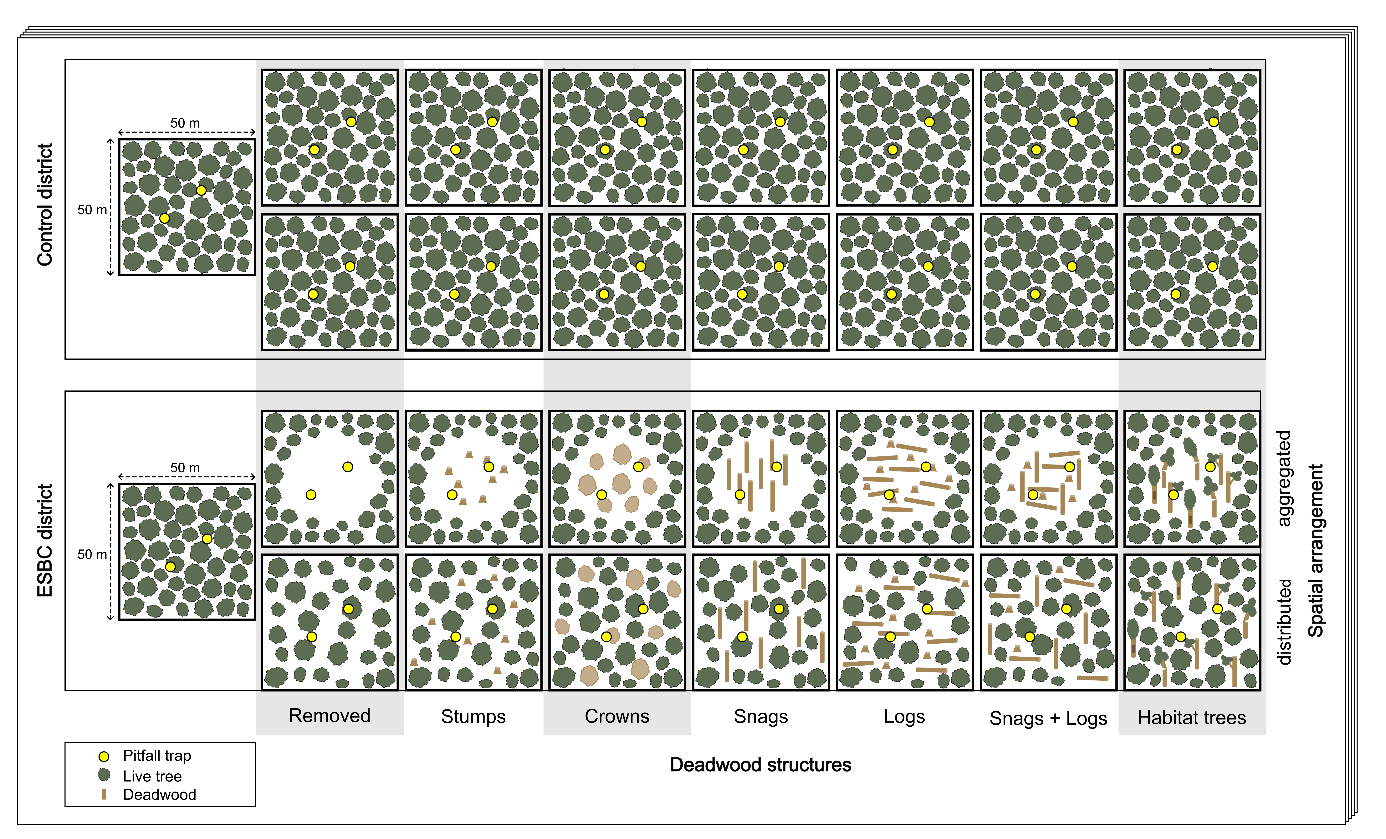

**Figure S1** Schematic illustration of an experimental forest site of the ΒETA-FOR research unit. One site consists of two different districts with the same number of patches. In the ESBC (Enhancing Structural Βeta Complexity) district, manipulations were performed to increase forest heterogeneity by creating canopy gaps and deadwood structures. The control district consisted of homogeneous, untreated patches with a closed canopy. The manipulations in the ESBC district were either performed spatially aggregated in the plot center, creating a central canopy gap, or distributed throughout the patch to maintain the closed canopy. The manipulations either removed the manipulated trees completely or created different types of deadwood structures: stumps, crowns, snags, logs, snags + logs, and habitat trees. Grey-colored treatments were applied only in the University Forest region. Two pitfall traps were installed at the center of each patch to assess spider diversity.
