## Supplementary Text for "Temperate forest heterogeneity decreases local and landscape-scale spider diversity through habitat filtering despite species turnover"

By intermixing functional and phylogeny distances through the weighting parameter *a*, a suitable MFPD in-between 0 for only functional distances and 1 for only phylogenetic distances is calculated, and with intermediate values both phylogenetic and functional information are contributing to MFPD. We used the function *funct.phylo.dist* from the *pez*-package (Pearse et al., 2015), the phylogenetic distance matrix was produced with the function *cophenetic* from the *stats* package (R Core Team, 2025) and the functional distance matrix was mentioned earlier. Functional-phylogenetic diversity was calculated as mean pairwise distances (MPD) based on the MFPD distance matrix. To control for variation in species numbers in the samples and obtain a metric on the assembly pattern, a null-model approach to MPD was applied using the tip-shuffling method (Cadotte & Davies, 2016). This results in standardized effect sizes (SES) of the mean pairwise distances (SES MPD), which were calculated with 999 randomizations with the function *ses.mpd* in the package *picante* (Kembel et al., 2010).

**References**

Cadotte, M., & Davies, J. (2016). *Phylogenies in Ecology: A Guide to Concepts and Methods*. https://doi.org/10.1515/9781400881192

Kembel, S. W., Cowan, P. D., Helmus, M. R., Cornwell, W. K., Morlon, H., Ackerly, D. D., Blomberg, S. P., & Webb, C. O. (2010). Picante: R tools for integrating phylogenies and ecology. *Bioinformatics*, *26*(11), 1463–1464. https://doi.org/10.1093/bioinformatics/btq166

Pearse, W. D., Cadotte, M. W., Cavender-Bares, J., Ives, A. R., Tucker, C. M., Walker, S. C., & Helmus, M. R. (2015). pez: phylogenetics for the environmental sciences. *Bioinformatics*, *31*(17), 2888–2890. https://doi.org/10.1093/bioinformatics/btv277

R Core Team. (2025). *R: A Language and Environment for Statistical Computing*. https://www.r-project.org/
