## Supplementary figures and images for "Temperate forest heterogeneity decreases local and landscape-scale spider diversity through habitat filtering despite species turnover"

### Fig. S2

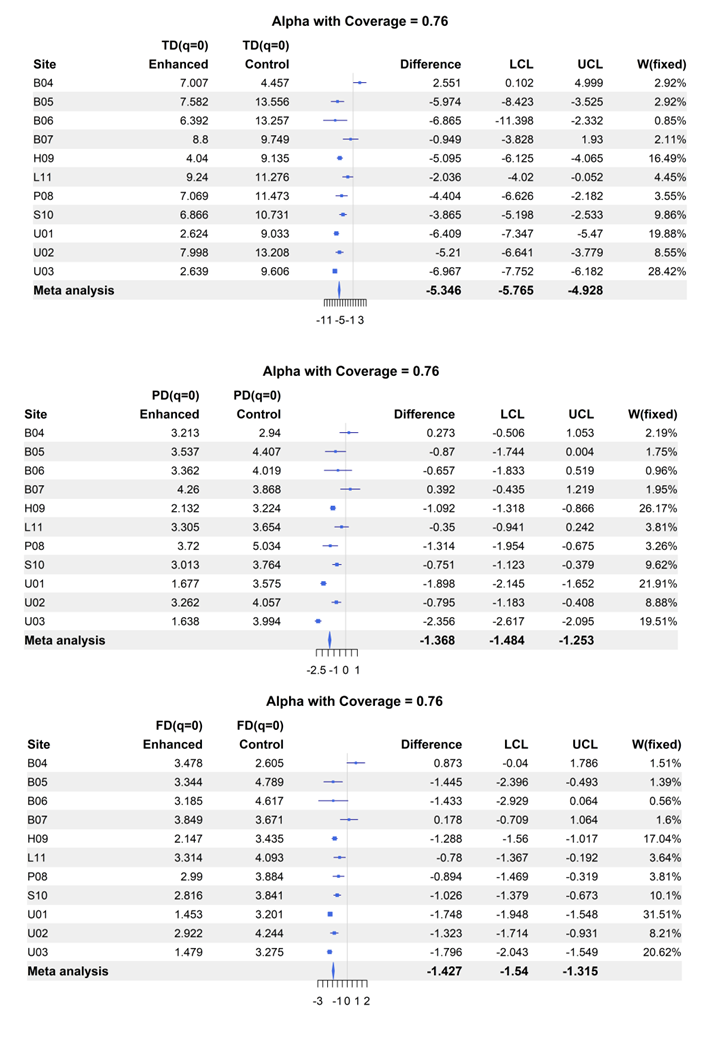
**Figure S2** Metaanalysis at the alpha level for q0.

### Fig. S3

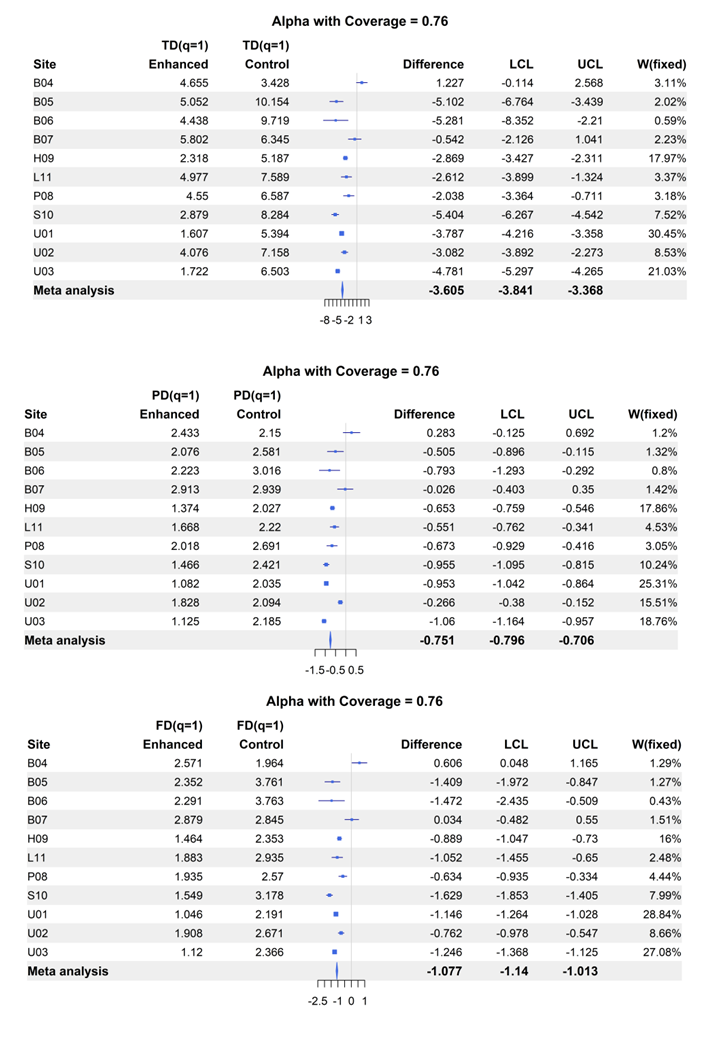
**Figure S3** Metaanalysis at the alpha level for q1.

### Fig. S4

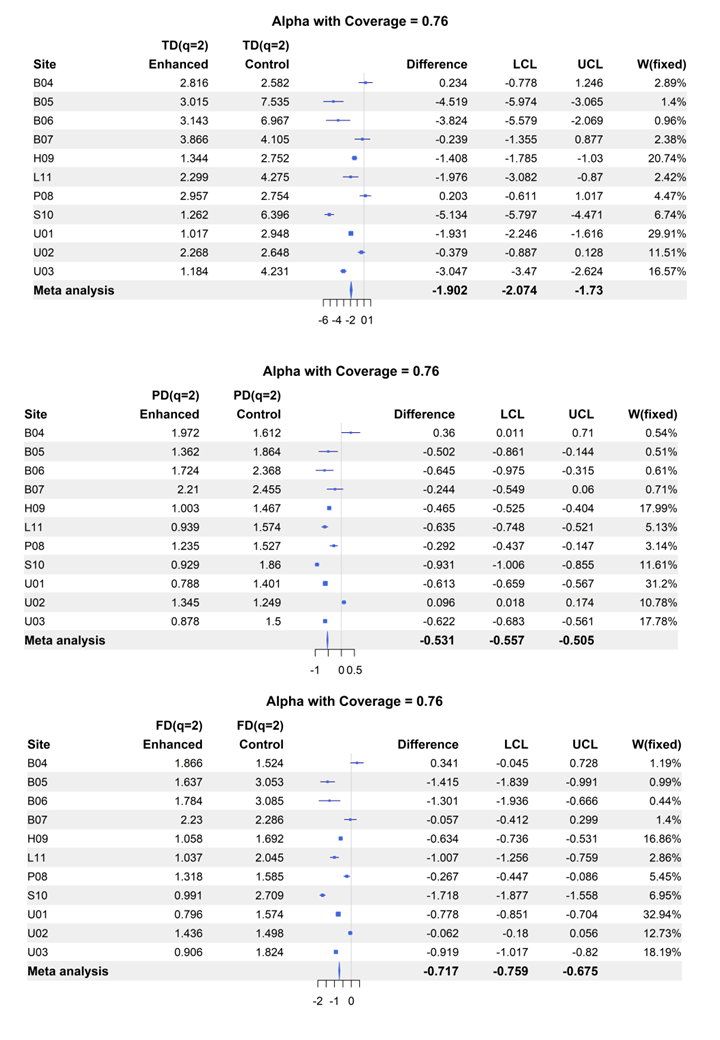
**Figure S4** Metaanalysis at the alpha level for q2

### Fig. S5

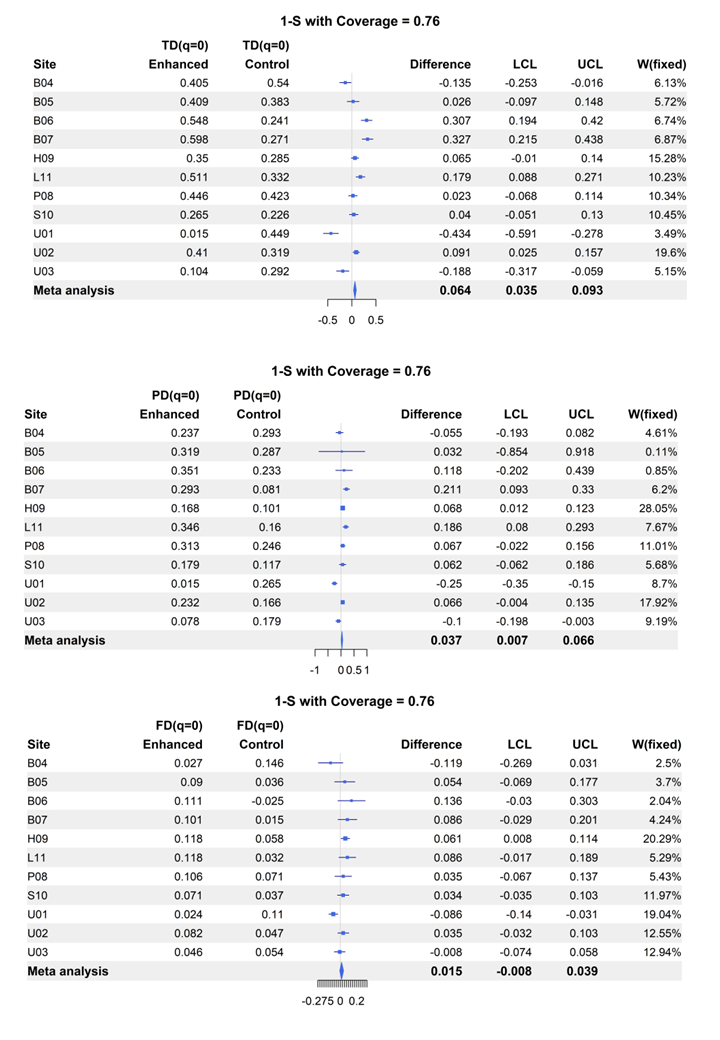
**Figure S5** Metaanalysis at the beta (1-S) level for q0.

### Fig. S6

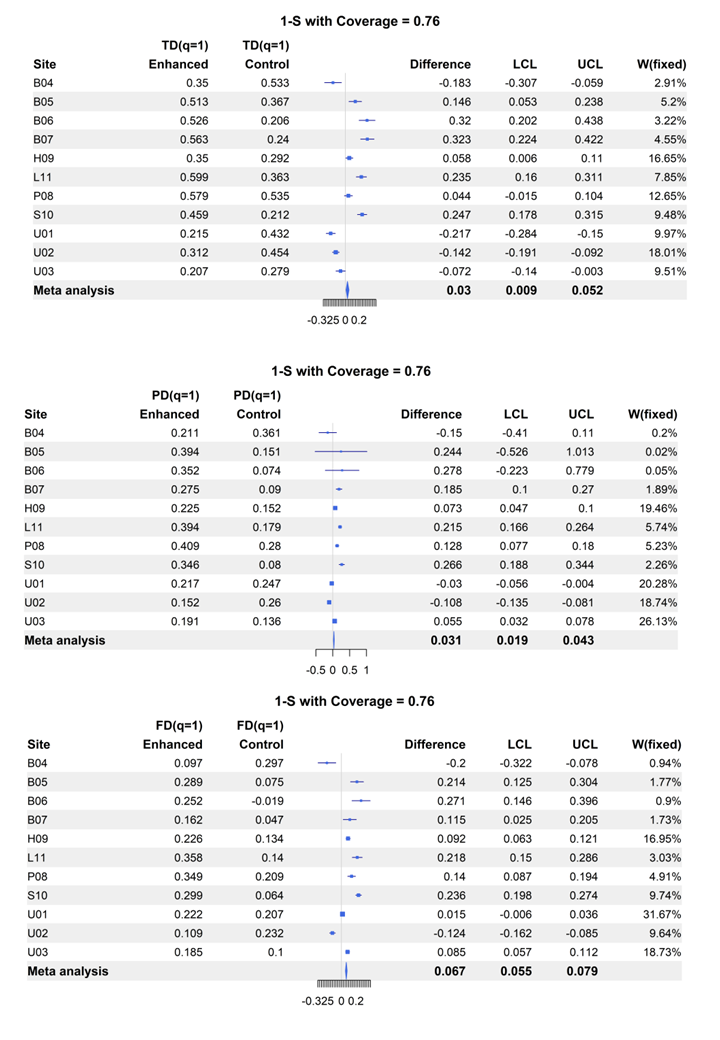
**Figure S6** Metaanalysis at the beta (1-S) level for q1.

### Fig. S7

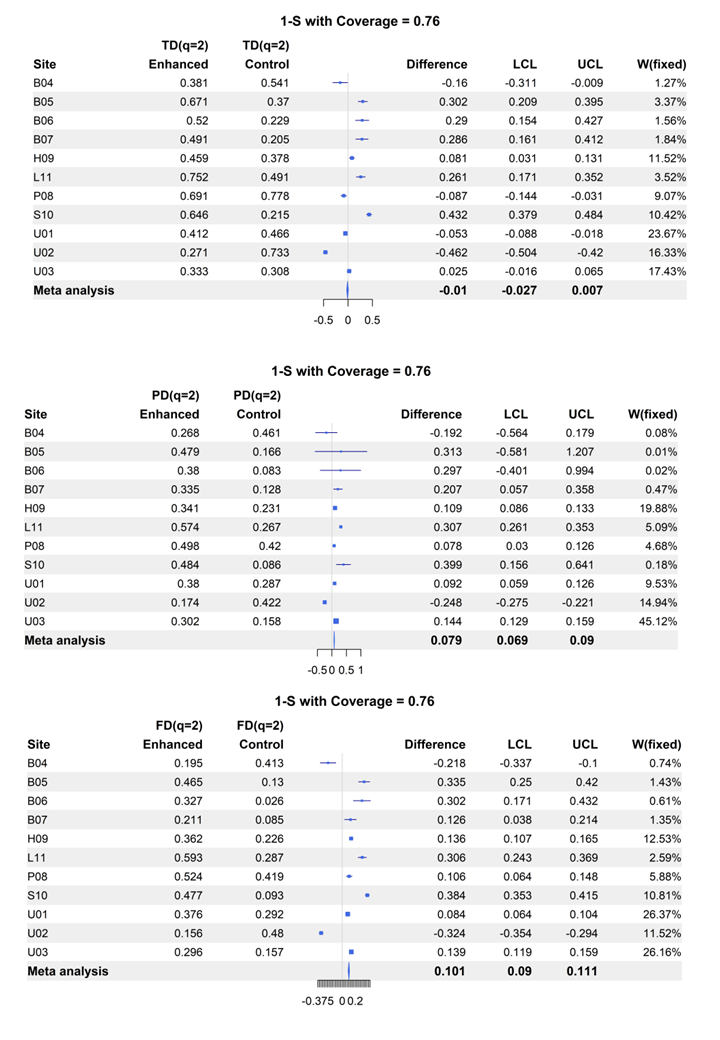
**Figure S7** Metaanalysis at the beta (1-S) level for q2.

### Fig. S8

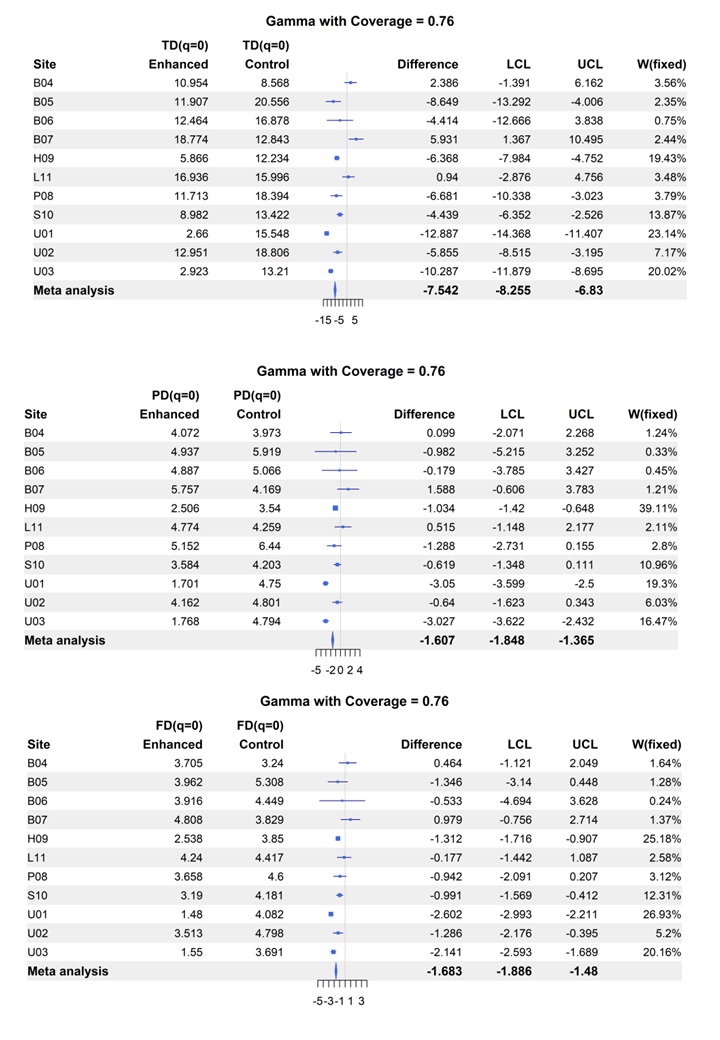
**Figure S8** Metaanalysis at the gamma level for q0.

### Fig. S9

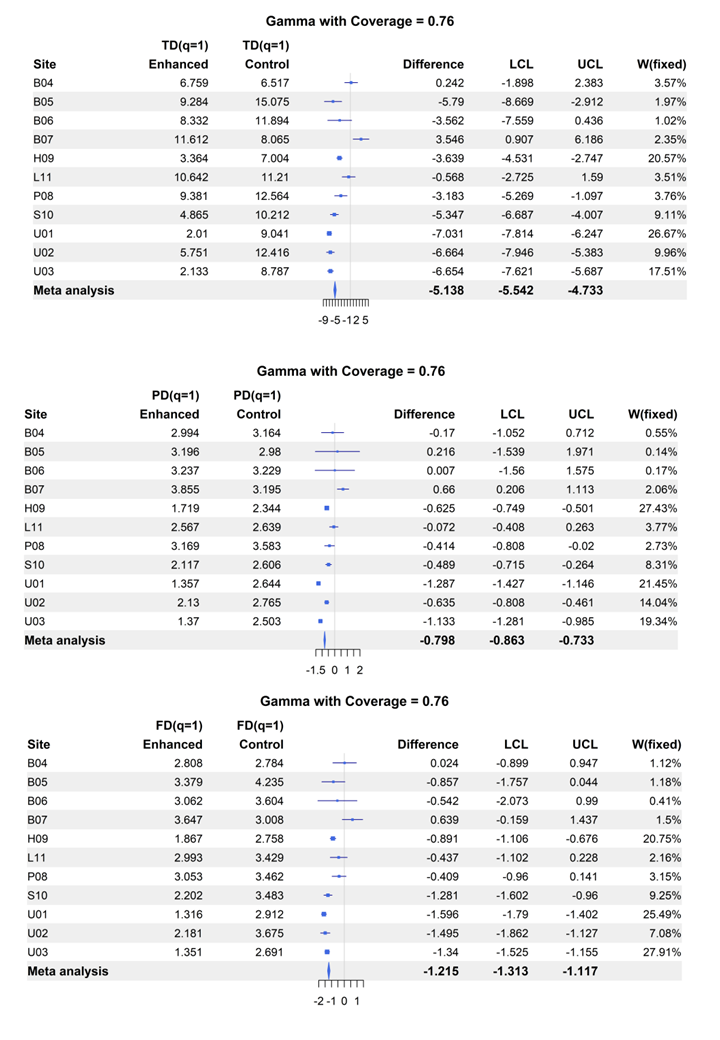
**Figure S9** Metaanalysis at the gamma level for q1.

### Fig. S10

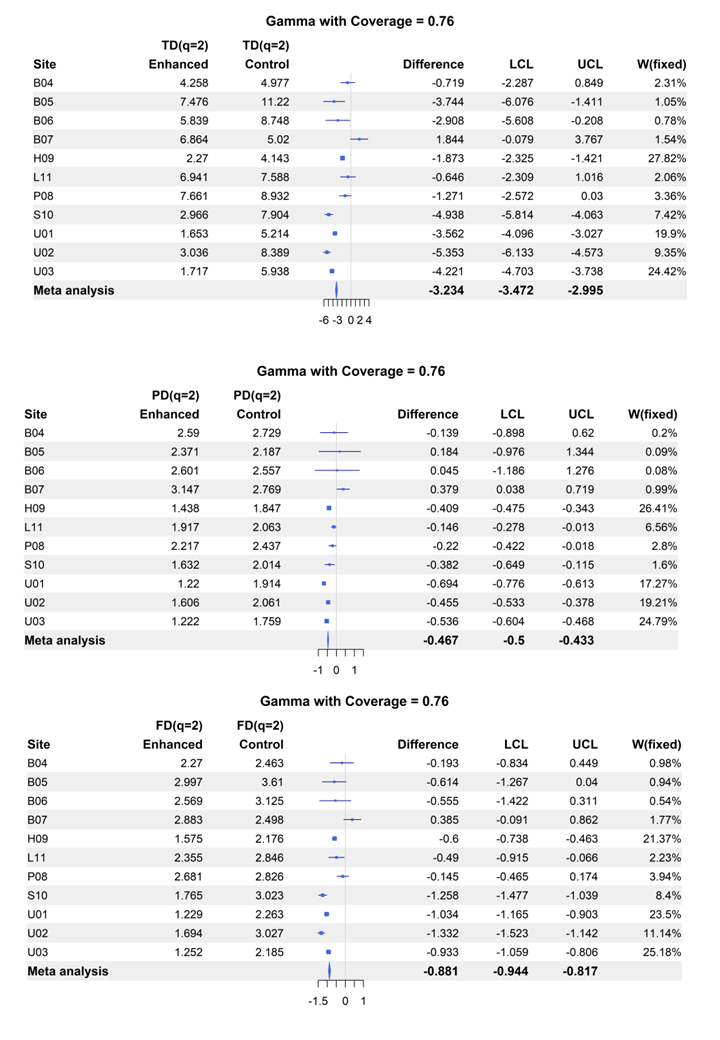
**Figure S10** Metaanalysis at the gamma level for q2.
