## Supplementary material for "Temperate forest heterogeneity decreases local and landscape-scale spider diversity through habitat filtering despite species turnover": Fig. S11

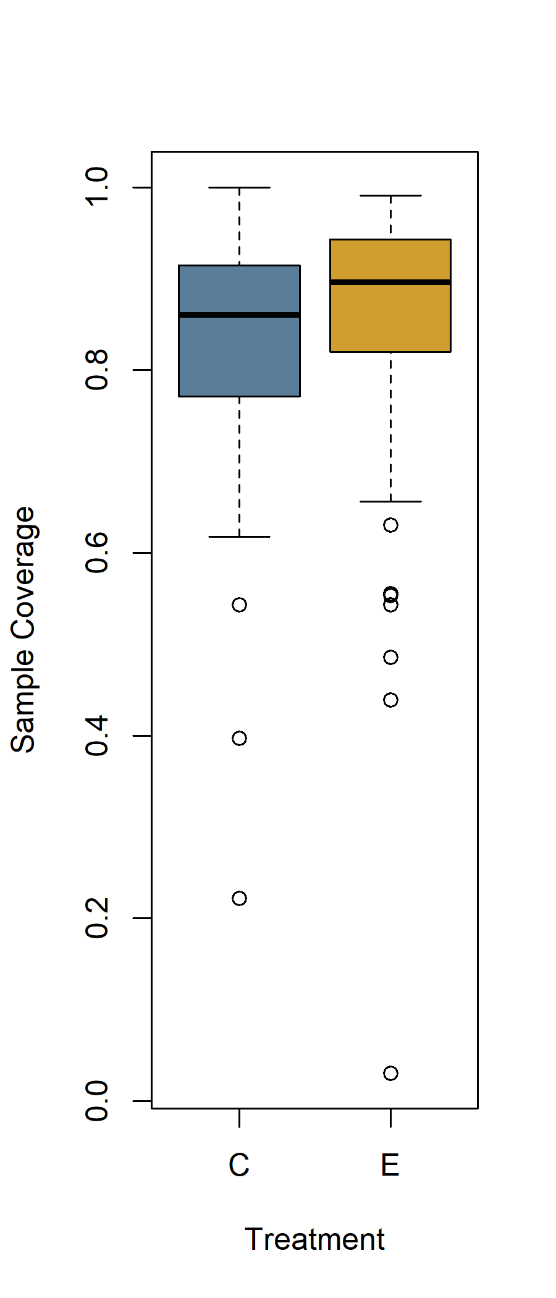

**Figure S11** Observed sample coverage of spider pitfall samples per patch (alpha-level). Sample coverage was higher on ESBC patches (treatment E) vs. control patches (treatment C), indicating a more complete sampling of spider diversity in heterogeneous, more open forests with pitfall traps.
