## Supplementary material for "Temperate forest heterogeneity decreases local and landscape-scale spider diversity through habitat filtering despite species turnover": Fig. S12

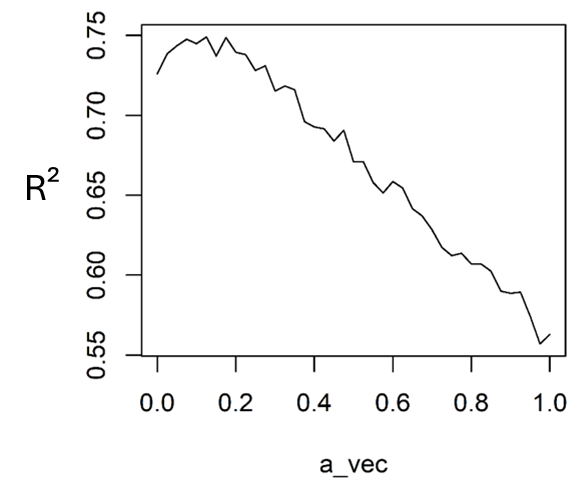

**Figure S12** Optimizing SES MFPD (nullmodel using label shuffling) along an increasing a-value, which gives an increasing weight to phylogenetic distance from 0 only functional to 100% only phylogeny. Based on the highest R² at 12% phylogeny the final functional diversity was calculated.
